# Morpho-anatomical observations on Phyllanthus of Southwestern Bangladesh with two new records for Bangladesh

**DOI:** 10.1101/608711

**Authors:** Md. Sajjad Hossain Tuhin, Md. Sharif Hasan Limon

**Affiliations:** Forestry and Wood Technology Discipline, Khulna University, Khulna 9208, Bangladesh

**Keywords:** Angiosperm, Phyllanthus, taxonomy, distribution, Bangladesh, *Phyllanthus amarus*, *Phyllanthus debilis*

## Abstract

An extensive floristic survey was done to annotate Phyllanthus of southwestern Bangladesh from 2015 to 2018. In total, 2189 individuals of Phyllanthus were counted and identified as eight different species (five herbs, two trees and a shrub). All species were examined following both morphological and anatomical methods, based on taxonomic notes. The listed species were *Phyllanthus acidus, Phyllanthus amarus, Phyllanthus debilis, Phyllanthus emblica, Phyllanthus niruri, Phyllanthus urinaria, Phyllanthus reticulatus* and *Phyllanthus virgatus*. Among them, *Phyllanthus amarus* and *Phyllanthus debilis* were listed for the first time from Bangladesh during this study period.

## Introduction

Phyllantheace is a very common taxa found in Bangladesh. It is a large family of flowering plants, consisting of 59 accepted genera, 10 tribes, two subfamilies and about 2000 species worldwide (Hoffmann et. al. 2006). This family is the second largest segregation of Euphorbiaceae sensu lato (s.l.) among Pandaceae, Picrodendraceae, and Putranjivaceae (Savolainen et al. 2000; Samuel et al. 2005). Phyllanthus is the largest genera of Phyllanthaceae consisting approximately 1000 species. Most of the Phyllanthus species are distributed in tropical and subtropical region (Sarin et al. 2014; Calixto et al. 1998). Previously, Phyllanthus represented 11 species (5 herbs, 4 shrubs and 2 trees) in Bangladesh (Ahamed et al. 2008). Most of the regional literatures of this region ensured a valuable contribution of it to the regional flora (Uddin & Hassan 2010; Islam et al. 2009; Rahman et. al. 2015; Tutul et. al. 2010). In addition to that, most of them are known to have economic and medicinal uses. Some of them had already been used in ethno-medicine and extraction of antioxidant and other industrial chemicals (Ali et al. 2006; Cesari et al. 2015; Gebhardt et al. 2003; Okafor et al. 2008; Patel et al. 2011; Thyagarajan et al. 1988). So, it is very important to conserve those species in terms of ecology and economy. But, many of the Phyllanthus sp. have a close morphological characteristics, which made them very difficult to differentiate at the species level (Kandavel et al. 2011; Webster 1956). Meanwhile, conservation demands proper identification to study its ecology and distribution. In this current study, available Phyllanthus individuals were collected from the south-western part of Bangladesh and attempted to identify at species level with morphological and anatomical features.

## Materials and Methods

### Study Site

Khulna division lies between 21.643° N to 24.181° N and 88.56° E to 89.943° E (Fig. 1). It experiences a tropical climatic condition with a mild winter from October to March, hot and humid summer in March to June and humid, warm rainy monsoon in June to October. Temperature varies all the year round, in January and December temperature falls to the lowest 12-15°C and it reached highest in April-June at 41-45 °C. Daily relative humidity lied between 50-90%, which became lowest in evening and highest in morning. Rainy season occurred here from June to October. Maximum precipitation was experienced in July with 20-25 days of rain with 368 mm precipitation. Wind speed is also variable here. Wind speed shows higher in the months of April, May, June July and August. In those months average wind speed is over 5 Km/h. In the month of May the average wind speed reaches height around 8 km/h. Maximum wind speed also reaches higher in this month is around 80-90 km/h. December is the calmest month of the year, wind speed is lowest here. Average daily wind speed during this month is 2 km/h and maximum reaches only 26 km/h (Bangladesh Bureau of Statistics 2014).

**Fig 1:**
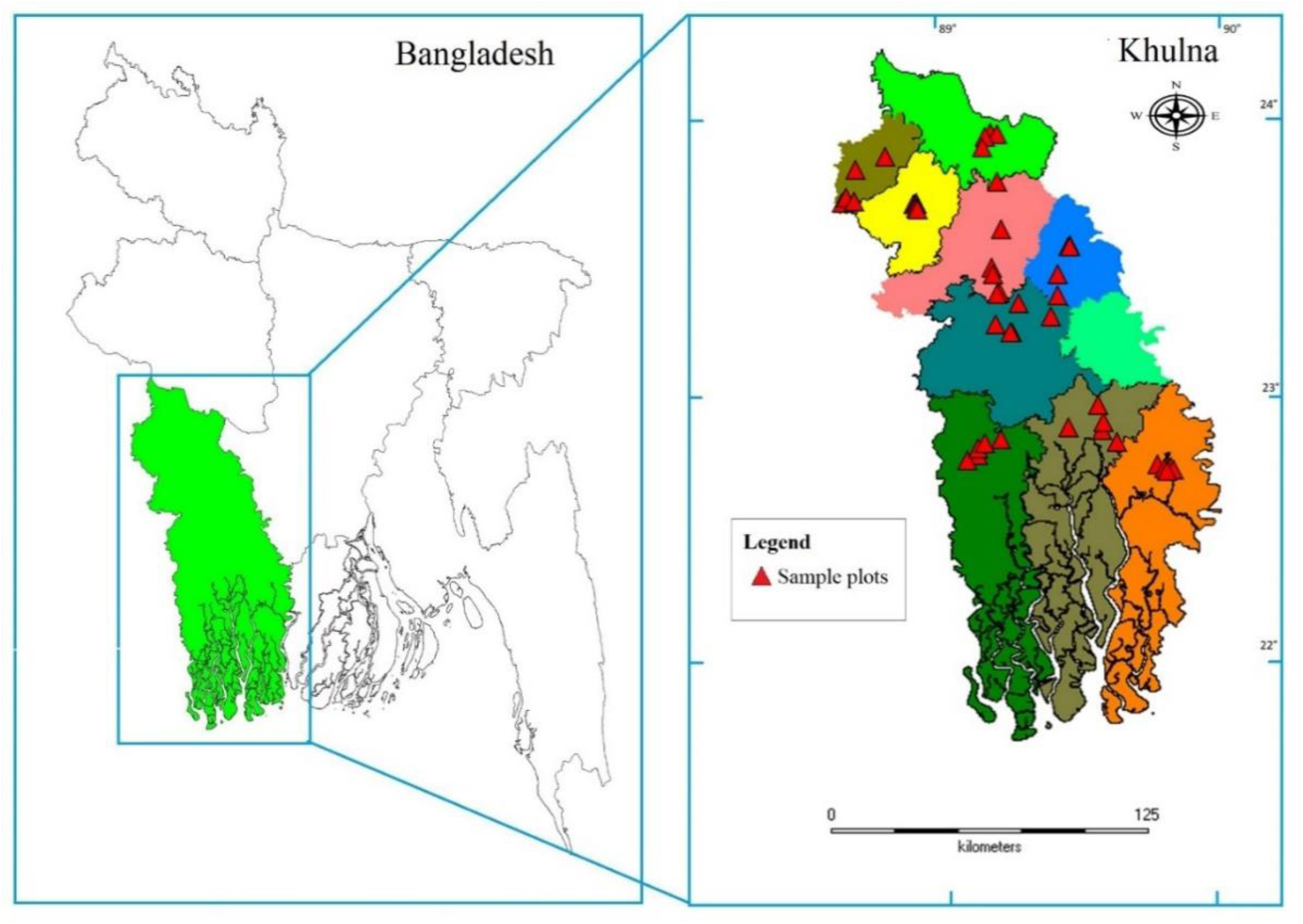
Map of the study area

### Sampling design

The study was carried out at nine administrative districts (namely, Khulna, Kushita, Jessore, Bagherhat, Chuadanga, Satkhira, Magura, Meherpur and Jhinaidha) of southwestern of Bangladesh from April 2015 to August 2018. In total, 45 sites were selected from those 9 districts (5 sites from each) by opportunistic sampling technique (Fig. 01, table 1) (Milchunas et al. 1992). Habitat, diversity of Phullanthus and population size were considered to select the study sites. Standard herbarium technique (Hyland 1972; Rahman et al. 2013) was used to collect the herbarium specimen for analysis. Live specimens were photographed by Celestron handheld digital microscope and Nikon-3200D camera with Nikkor 18-55mm AF-S DX, Nikkor 55-300mm AF-S DX and Micro-Nikkor 105 mm FX AF lenses. We recorded GPS quadrate for all surveyed sites with a GPS device (Garmin GPSMAP 76CSx). Image J (1.52a) was used to record morphological measurements and image analysis. Herbarium sheets were prepared and deposited to FWT herbarium to conserve for future use. We did not collect any endangered angiosperm or sample from any protected areas, hence no permission was required for the collections. Collected specimens were analyzed and identified based on key provided by Hooker (1890), Prain (1904), Kanjilal et al. (1938), Deb (1983), Matthew (1999) and Ahmed et. al. (2008).

**Table 1:** A list of locations in the southwestern region of Bangladesh surveyed during the study period

| Sl no. | District | Place | Latitude | Longitude | No. of individuals | No. of species |
| --- | --- | --- | --- | --- | --- | --- |
| 1. | Khulna | KU Campus | 22.80125° N | 89.53466° E | 117 | 6 |
| 2. |  | Boyra | 22.83289° N | 89.54131° E | 46 | 5 |
| 3. |  | Dumuria | 22.80936° N | 89.41360° E | 67 | 5 |
| 4. |  | Dhigholia | 22.89416° N | 89.52221° E | 32 | 4 |
| 5. |  | Baintala | 22.75812° N | 89.58919° E | 13 | 3 |
| 6. | Jessore | Pouro park | 23.16508° N | 89.20466° E | 41 | 4 |
| 7. |  | M. M. collage | 23.16234° N | 89.20328° E | 33 | 4 |
| 8. |  | Cantonment | 23.19148° N | 89.14975° E | 41 | 6 |
| 9. |  | Labutala | 23.27200° N | 89.23148° E | 49 | 6 |
| 10. |  | Bagherpara | 23.22126° N | 89.34825° E | 11 | 4 |
| 11. | Satkhira | Govet. collage | 22.71028° N | 89.08503° E | 23 | 5 |
| 12. |  | Medical collage | 22.69100° N | 89.04764° E | 41 | 5 |
| 13. |  | BGB camp | 22.73244° N | 89.08453° E | 78 | 5 |
| 14. |  | Binerpota | 22.75447° N | 89.10717° E | 47 | 4 |
| 15. |  | Patkelgatha | 22.76752° N | 89.16900° E | 39 | 4 |
| 16. | Bhagerhat | Satgomuj Mosque | 22.67461° N | 89.73788° E | 71 | 4 |
| 17. |  | Khanjahan Ali<br>Majar | 22.66067° N | 89.76032° E | 23 | 3 |
| 18. |  | PC College | 22.66543° N | 89.78261° E | 87 | 6 |
| 19. |  | Pouro Lake | 22.65560° N | 89.79535° E | 21 | 4 |
| 20. |  | Pocha Dighi | 22.64870° N | 89.77262° E | 23 | 5 |
| 21. | Magura | Govt. HSS<br>College | 23.48938° N | 89.41914° E | 19 | 3 |
| 22. |  | Stadium Para | 23.48232° N | 89.41420° E | 40 | 4 |
| 23. |  | District Fishery<br>Office | 23.48080° N | 89.41904° E | 47 | 5 |
| 24. |  | Arpara | 23.37775° N | 89.37197° E | 38 | 5 |
| 25. |  | Salikha | 23.30223° N | 89.37575° E | 112 | 6 |
| 26. | Jhinaidha | KC College | 23.54539° N | 89.17012° E | 13 | 2 |
| 27. |  | Kaligang | 23.40495° N | 89.13213° E | 35 | 4 |
| 28. |  | Dulal-Mundia | 23.38004° N | 89.13924° E | 72 | 5 |
| 29. |  | Gazi-Kalu<br>Mazar | 23.30836° N | 89.16321° E | 67 | 6 |
| 30. |  | Barobazar<br>Railway Station | 23.30488° N | 89.15081° E | 93 | 6 |
| 31. | Kushtia | Kushtia Govt.<br>College | 23.90175° N | 89.12837° E | 73 | 6 |
| 32. |  | Kushtia Soshan | 23.89993° N | 89.15211° E | 65 | 5 |
| 33. |  | Chourhas | 23.88921° N | 89.10866° E | 34 | 5 |
| **34.** |  | Bot-toil | 23.85114° N | 89.09854° E | 59 | 5 |
| **35.** |  | IU Campus | 23.72105° N | 89.15204° E | 20 | 6 |
| **36.** | Chuadanga | Chuadanga<br>Govt. College | 23.63979° N | 88.84907° E | 67 | 5 |
| **37.** |  | Rail Para | 23.64256° N | 88.85871° E | 17 | 4 |
| **38.** |  | Bus Terminal | 23.63715° N | 88.86063° E | 90 | 5 |
| **39.** |  | Jafarpur | 23.62353° N | 88.85984° E | 42 | 5 |
| **40.** |  | BGB Camp | 23.62049° N | 88.86415° E | 65 | 5 |
| **41.** | Meherpur | Mujibnagar | 23.64411° N | 88.59134° E | 43 | 5 |
| **42.** |  | Kedergong | 23.66181° N | 88.60596° E | 34 | 3 |
| **43.** |  | Bollovpur | 23.64793° N | 88.63281° E | 51 | 3 |
| **44.** |  | Collage road | 23.76746° N | 88.63974° E | 9 | 2 |
| **45.** |  | Gangni | 23.81729° N | 88.74912° E | 81 | 5 |

## Result

2189 individuals of Phyllanthus were examined and classified into eight different species from southwestern part of Bangladesh. Among these, five were herbs, two trees and a shrubs. Expect *Phyllanthus debilis* all the species were well distributed throughout the study area. Meanwhile, *Phyllanthus niruri, Phyllanthus urinaria, Phyllanthus amarus* and *Phyllanthus reticulatus* were very abundant in the study sites. But *Phyllanthus virgatus* were found in Jessore-Chuadangha belt and absent in Khulna, Satkhira and Bhagerhat (table 2). Apart from these, *Phyllanthus acidus* and *Phyllanthus emblica* could be listed as a plantation species as they were absent at natural habitats. In addition to it, *Phyllanthus amarus* and *Phyllanthus debilis* were listed for the first time from Bangladesh at current study.

**Table 2:**
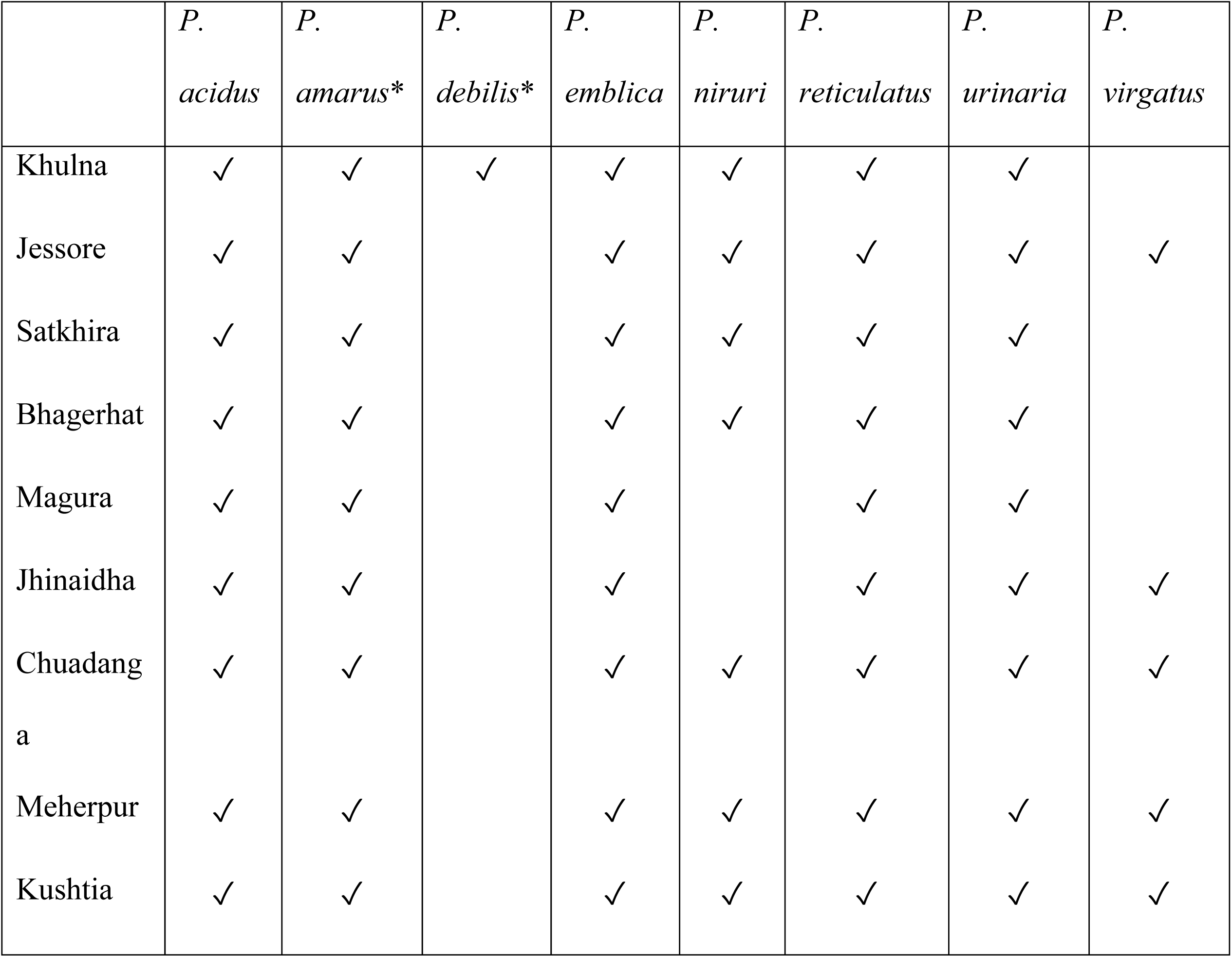
List and distribution of the listed *Phyllanthus sp.* at current study (* new record for Bangladesh).

### *Phyllanthus acidus* (L.) Skeels: In: Bull. Bur. Pl. Industr. U.S.D.A. 148: 17 (1909)

#### Synonyms

*Averrhoa acida* L.: In: Sp. Pl.: 428 (1753), *Cicca acida* (L.) Merr.: In: Interpr. Herb. Amboin.: 314 (1917), *Cicca acidissima* Blanco: In: Fl. Filip.: 700 (1873), *Cicca disticha* L.: In: Mant. Pl. 1: 124 (1767), *Cicca nodiflora* Lam.: In: Encycl. 2: 1 (1786), *Cicca racemosa* Lour.: In: Fl. Cochinch.: 556 (1790), *Diasperus acidissimus* (Blanco) Kuntze: In: Revis. Gen. Pl. 2: 598 (1891), *Phyllanthus acidissimus* (Blanco) Müll. Arg.: In: Linnaea 32: 50 (1863), *Phyllanthus cicca* Müll. Arg.: In: Linnaea 32: 50 (1863), *Phyllanthus cicca* var. bracteosa Müll.Arg.: In: Linnaea 32: 50 (1863), *Phyllanthus cochinchinensis* (Lour.) Müll.Arg., nom. illeg.: In: Prodr. 15(2): 417 (1866), *Phyllanthus distichus* (L.) Müll.Arg., nom. illeg.: In: Prodr. 15(2): 413 (1866), *Phyllanthus distichus* f. nodiflorus (Lam.) Müll.Arg.: In: Prodr. 15(2): 414 (1866), *Phyllanthus longifolius* Jacq.: In: Pl. Hort. Schoenbr. 2: 36 (1797), *Tricarium cochinchinense* Lour.: In: Fl. Cochinch.: 557 (1790).

A Monoecious, glabrous, deciduous, small sized tree, up to 15m high; branches stout, leafless, with slender deciduous leafy branchlets at the end; bark brownish. Leaves distichous, alternate, stipulate, stipules lanceolate, 0.8-1 mm; petiolate, 1.5-3mm long, distichous; leaf blade ovate to ovate-lanceolate, 3-8mm long and 1-3.5mm wide, leathery, acute to acuminate at apex, rounded or broadly cuneate at base, lateral veins 7-9 pairs, sometimes indistinct. Flower bisexual, dark reddish, at dense clusters forming slender, glabrous, in interrupted racemes, arising along the stem and branches. Male flowers are minute, red; staminate, stamen 4, free, 2 shorter; disc gland 4, smooth, diameter 0.4-0.7mm; pedicellate, pedicels 1-3 mm long and less than 1 mm width, slender; sepal 4, suborbicular, 0.8-1mm long, entire. Female flowers are very few, pedicellate; sepal 4, 1-1.5 mm long, apex rounded; ovary glabrous, 3-4 celled; styles 4, free, fleshy, bifid, recurved. Fruit a drupe, depressed globose to subglobose, smooth, 1.5-2.5 cm in diameter, 6-8 lobed, exocarp fleshy, pale yellowish green, endocarp crustaceous. Seeds reddish brown, trigonous or convex, 2-4mm long. Flowering and fruiting: March-July (Fig. 2).

**Fig. 2:**
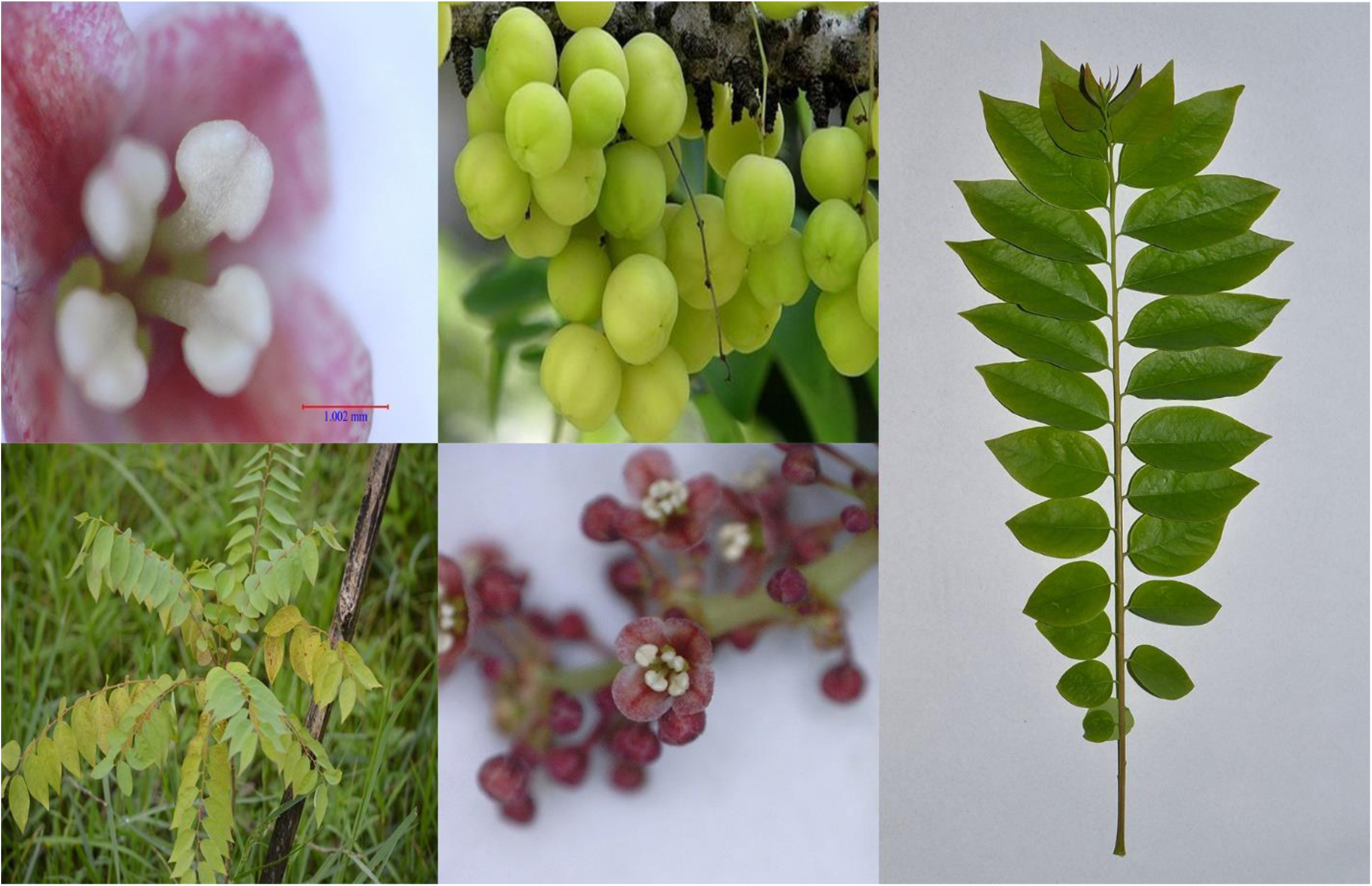
Different parts of *Phyllanthus acidus* (L.) Skeels

### *Phyllanthus amarus* Schumach. & Thonn.: In: Beskr. Guin. Pl.: 421 (1827)

#### Synonyms

*Phyllanthus swartzii* Kostel.: In: Allg. Med.-Pharm. Fl. 5: 1771 (1836), *Phyllanthus niruroides* var. *madagascariensis* Leandri, nom. inval.: In: Notul. Syst. (Paris) 7: 184 (1839), *Phyllanthus scabrellus* Webb: In: Niger Fl.: 175 (1849), *Phyllanthus nanus* Hook.f.: In: Fl. Brit. India 5: 298 (1887), *Diasperus nanus* (Hook.f.) Kuntze: In: Revis. Gen. Pl. 2: 601 (1891), *Phyllanthus amarus* var. *baronianus* Leandri, nom. inval.: In: Notul. Syst. (Paris) 7: 183 (1939), *Phyllanthus niruri* var. *amarus* (Schumach. & Thonn.) Leandri: In: Fl. Madag. 111: 73 (1958), *Phyllanthus niruri* var. *baronianus* (Leandri) Leandri, nom. inval.: In: Fl. Madag. 111: 74 (1958), *Phyllanthus niruri* var. *scabrellus* (Webb) Müll.Arg.: In: Linnaea 32: 43 (1963).

A Monoecious erect annual herb, up to 1m high; stems often branching phyllanthoid with age, branches angular, 3-10 cm long (can reach up to 15 cm), usually with 20-42 leaves. Leaves alternate, membranous, oblong or elliptic-oblong; 3-8 mm long and 2-5mm wide, membranous or thinly papery, base rounded, apex obtuse or rounded and often apiculate; stipulate, lanceolate, scarious, acute, 0.8-1m long; petiolate, petioles very short 0.2-0.7mm long; entire leaf margin, lateral nerves 4-7 pairs, slightly conspicuous abaxially; indistinct, dark green above, paler and greyish beneath. Flower bisexual, light yellow, very numerous, axillary, along lower part of leafy shoots usually male, those in middle usually often bisexual with 1 female and 1 male flower, those toward branchlet apex often female. Male flowers are staminate, stamen 3, filamentous, filaments connate; pedicellate, pedicels 0.5-1 mm long and less than 0.4 mm width; sepal 5, 0.2-0.5 mm long, elliptic or obovate, apex acute; midrib yellowish green, margin membranous; disc gland 5, lobulated, orbicular or obovate, filaments completely connate into a column, 0.2-0.3 mm high; anthers sessile, 1 often reduced to a single anther sac (or sometimes only 2 functional anthers present), anther sacs divergent, slits completely confluent, horizontal. Female flower are pedicellate, pedicle 0.5-1.0 mm long; sepal 5, obovate to obovate-oblong, margin membranous, 0.8-1 mm long and 0.4-0.5 mm width, apex obtuse or acute; lobulate, 5 lobed, nectary disk thin and flat or subulate; ovary globose-triangular, 0.5 × 0.6 mm, smooth; styles free, erect or ascending, apex shallowly bifid. Fruit a capsule, depressed-globose, approximately 1-1.6 mm in diameter, dilated at apex, the surface smooth, explosively dehiscent. Seeds sharply 3-angled, 0.8-1 mm long and 0.6-0.8 mm width, light brown or yellowish brown, longitudinally 5-6 ribs, minutely transversely striate with hygroscopic cells; Flowering and fruiting: July-October (Fig. 3).

**Fig. 3:**
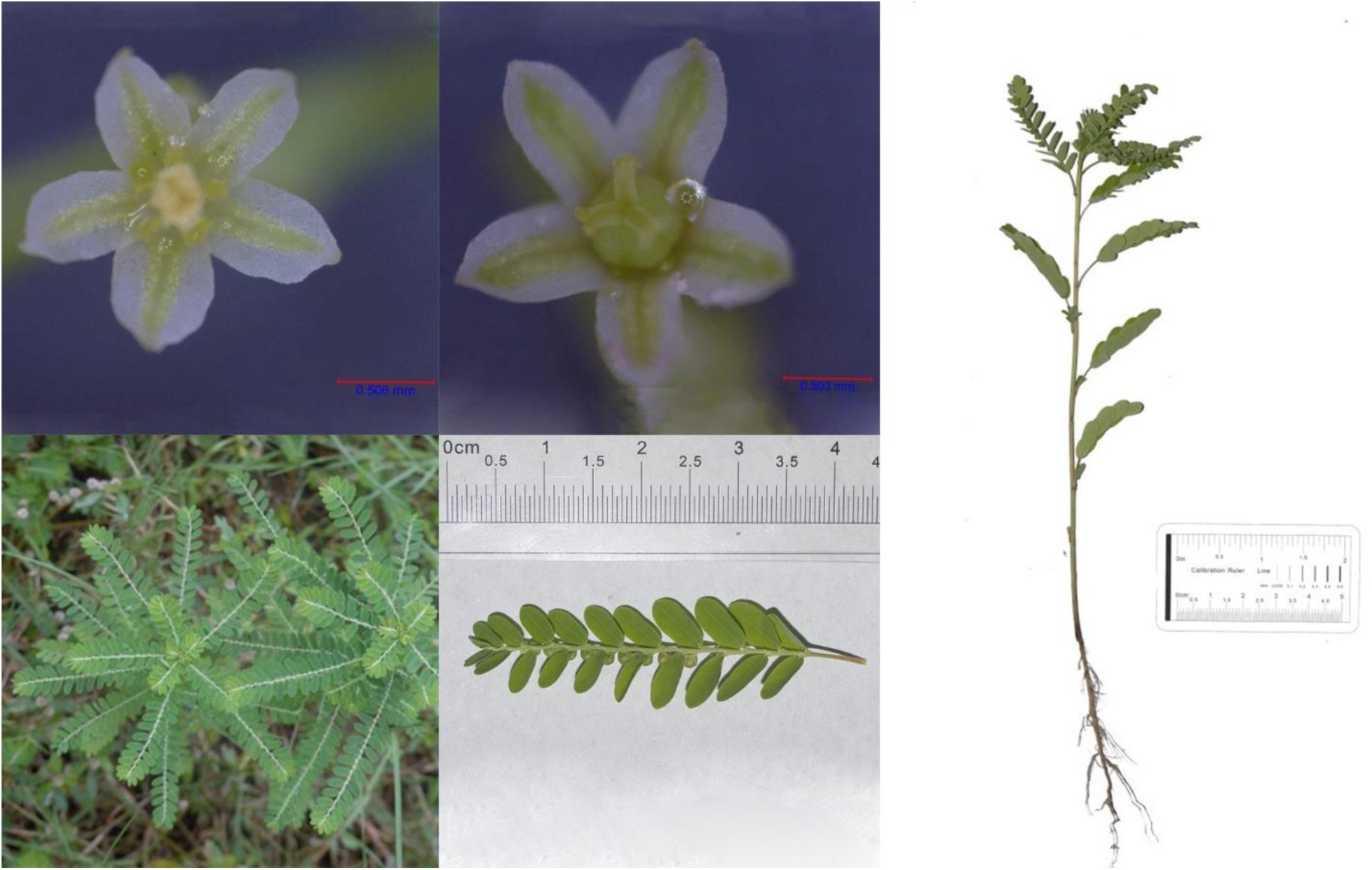
Different parts of *Phyllanthus amarus* Schumach. & Thonn

### *Phyllanthus debilis* J.G.Klein ex Willd.: In: Sp. Pl. 4: 582 (1805)

#### Synonyms

*Phyllanthus niruri* var. *javanicus* Müll.Arg.: In: Linnaea 32: 43 (1863), *Phyllanthus niruri* var. *debilis* (J.G.Klein ex Willd.) Müll.Arg.: In: Prodr. 15(2): 407 (1866), *Diasperus debilis* (J.G.Klein ex Willd.) Kuntze: In: Revis. Gen. Pl. 2: 601 (1891), *Phyllanthus boninsimae* Nakai: In: Bot. Mag. (Tokyo) 26: 96 (1912), *Phyllanthus leai* S.Moore: In: J. Linn. Soc., Bot. 45: 217 (1920).

A Monoecious erect annual herb, up to 60 cm high; stems often branching phyllanthoid with age, branches angular, 3-8 cm long (can reach up to 12 cm), usually with 12-35 leaves. Leaves alternate, stipulate, lanceolate, scarious, acute, 1.0-1.5mm long; petiolate, petioles very short 0.3-1mm long; narrowly elliptic to lanceolate, 8-20 mm long and 2-5mm wide, acute to acuminate at the apex and cuneate to acute at the base, entire leaf margin, membranous, lateral nerves 5-7 pairs, indistinct, dark green above, paler and greyish beneath. Flower bisexual, light yellow, very numerous, axillary. Male flowers 3-4 racemose cymes arising first 2-4 node of the branches; staminate, stamen 3, filamentous, filaments connate; pedicellate, pedicels 0.4-0.5 mm long and less than 0.5 mm width; sepal 6, 0.5-0.7 mm long, obovate, rounded, midrib yellowish green; disc gland 6, lobulate; anther horizontal. Female flower solitary in the node above male flowers; pedicellate, pedicle 1-1.6 mm long; sepal 6, obovate to oblong, rounded, 1.2-1.6 mm long and 0.4-0.6 mm width, lobulate, 6 lobed, margin whitish and scarious, nectary disk thin and flat, deeply lobed into 6-10, bifid; ovary subglobose, 0.8-1.1 mm diameter, smooth, styles minute and free. Fruit a capsule, depressed-globose, approximately 2-2.2 mm in diameter, the surface smooth, explosively dehiscent. Seeds angled ca. 1 mm long and 0.6 mm width, longitudinally 6-7 ribs and fine transverse striations on back, pale yellowish brown. Flowering and fruiting: March-June (Fig. 4).

**Fig. 4:**
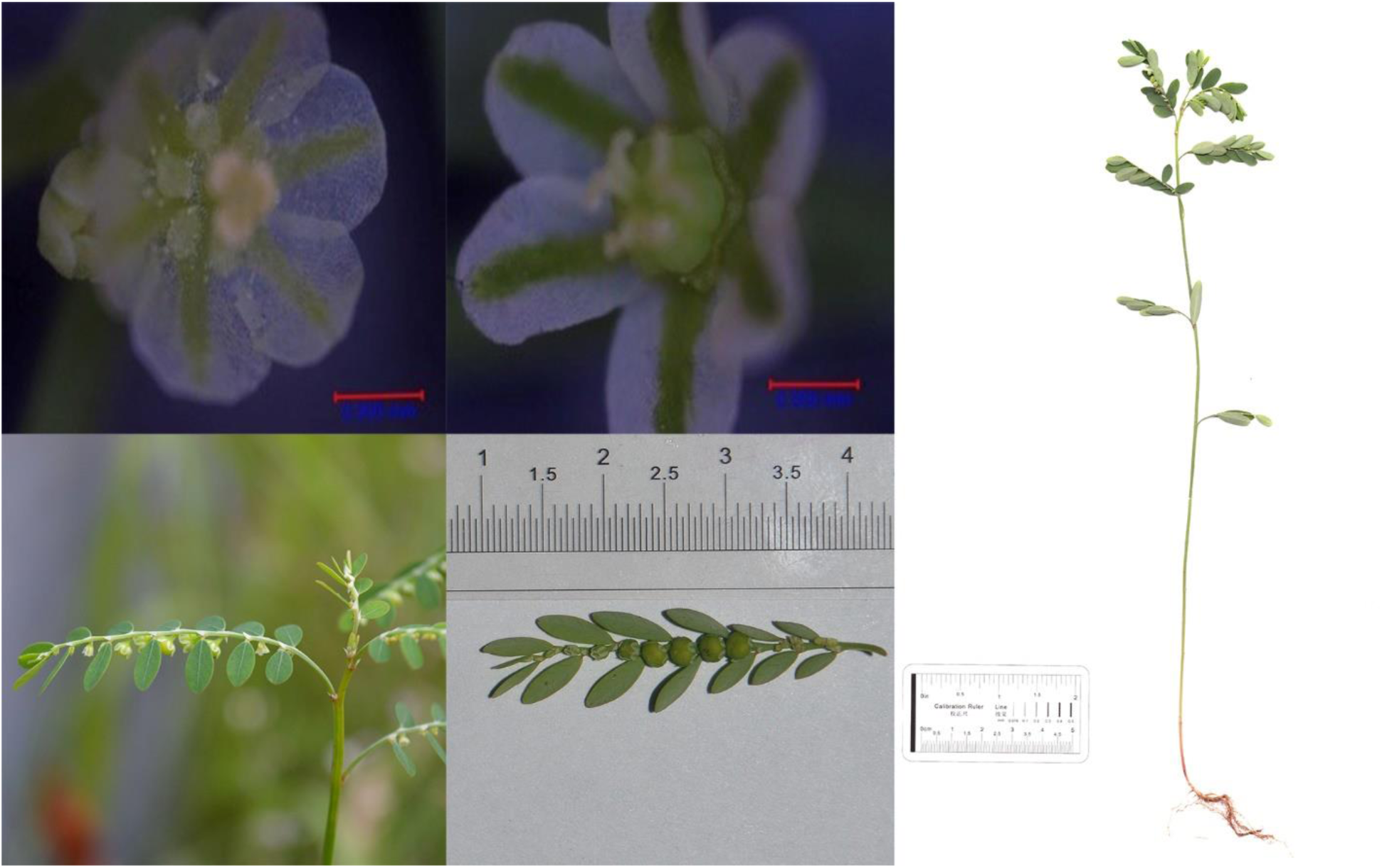
Different parts of *Phyllanthus debilis* J.G.Klein ex Willd.

### *Phyllanthus emblica* L.: In: Sp. Pl.: 982 (1753)

#### Synonyms

*Cicca emblica* (L.) Kurz: In: Forest Fl. Burma 2: 352 (1877), *Diasperus emblica* (L.) Kuntze: In: Revis. Gen. Pl. 2: 599 (1891), *Dichelactina nodicaulis* Hance: In: Ann. Bot. Syst. 3: 376 (1853), *Emblica arborea* Raf.: In: Sylva Tellur.: 91 (1838), *Emblica officinalis* Gaertn.: In: Fruct. Sem. Pl. 2: 122 (1970), *Phyllanthus glomeratus* Roxb. ex Benth., nom. nud.: In: Numer. List: n. ° 7903 (1847), *Phyllanthus mairei* H.Lév.: In: Bull. Acad. Int. Géogr. Bot. 25: 23 (1915), *Phyllanthus mimosifolius* Salisb.: In: Prodr. Stirp. Chap. Allerton: 391 (1796), *Phyllanthus taxifolius* D.Don: In: Prodr. Fl. Nepal.: 63 (1825).

A Monoecious, phuescent or glabrous, deciduous, medium sized tree, up to 25m high; bark brownish; main stems terete, sparsely lenticellate; leafy shoots angular, tawny pubescent. Leaves distichous, alternate, stipulate, stipules triangular-ovate, 0.8-1.5 mm; petiolate, petioles very short 0.3-0.8mm long, attenuate to acuminate, ciliolate, brown; leaf blade oblong or linear-oblong, 8-25mm long and 1.5-5mm wide, papery to leathery, paler abaxially, green adaxially, drying reddish or brownish, base shallowly cordate and slightly oblique, margin narrowly revolute, apex truncate, rounded or obtuse, mucronate or retuse at tip; lateral veins 4-9 pairs, sometimes indistinct. Flower bisexual, in axillary cymes, with many male flowers and sometimes 1 or 2 larger female flowers per cyme. Male flowers are staminate, stamen 3, filamentous, filaments coherent into a central column, 0.3-0.7mm long; anthers erect, obolong, 0.5-1mm long, horizontal; pedicellate, pedicels 1-3 mm long and less than 1 mm width, slender; sepal 6, 1.5-2.5 mm long and 0.5-1mm wide, obovate or spathulate, subequal, obtuse or rounded, entire, midrib yellowish green with a pale hyaline margin; disc gland 6, smooth, 0.2-0.5mm diameter. Female flowers are pedicellate, pedicle ca. 0.7mm long; sepal 6, oblong or spathulate, 1.5-2.5 mm long and 0.6-1.3 mm width, apex obtuse or rounded, thicker, margin membranous; ovary ovoid, ca. 1.5 mm, 3-celled; styles 3, stout, fleshy, 1.5-4 mm long, connate at base, apex deeply bifid, lobes divided at tip. Fruit a drupe, subglobose, smooth, 1-1.3 cm in diameter, exocarp fleshy, pale green or yellowish white, endocarp crustaceous. Seeds reddish, trigonous or plano-convex, 3-6mm long and 2-3 mm wide. Flowering and fruiting: March-October (Fig. 5)

**Fig. 5:**
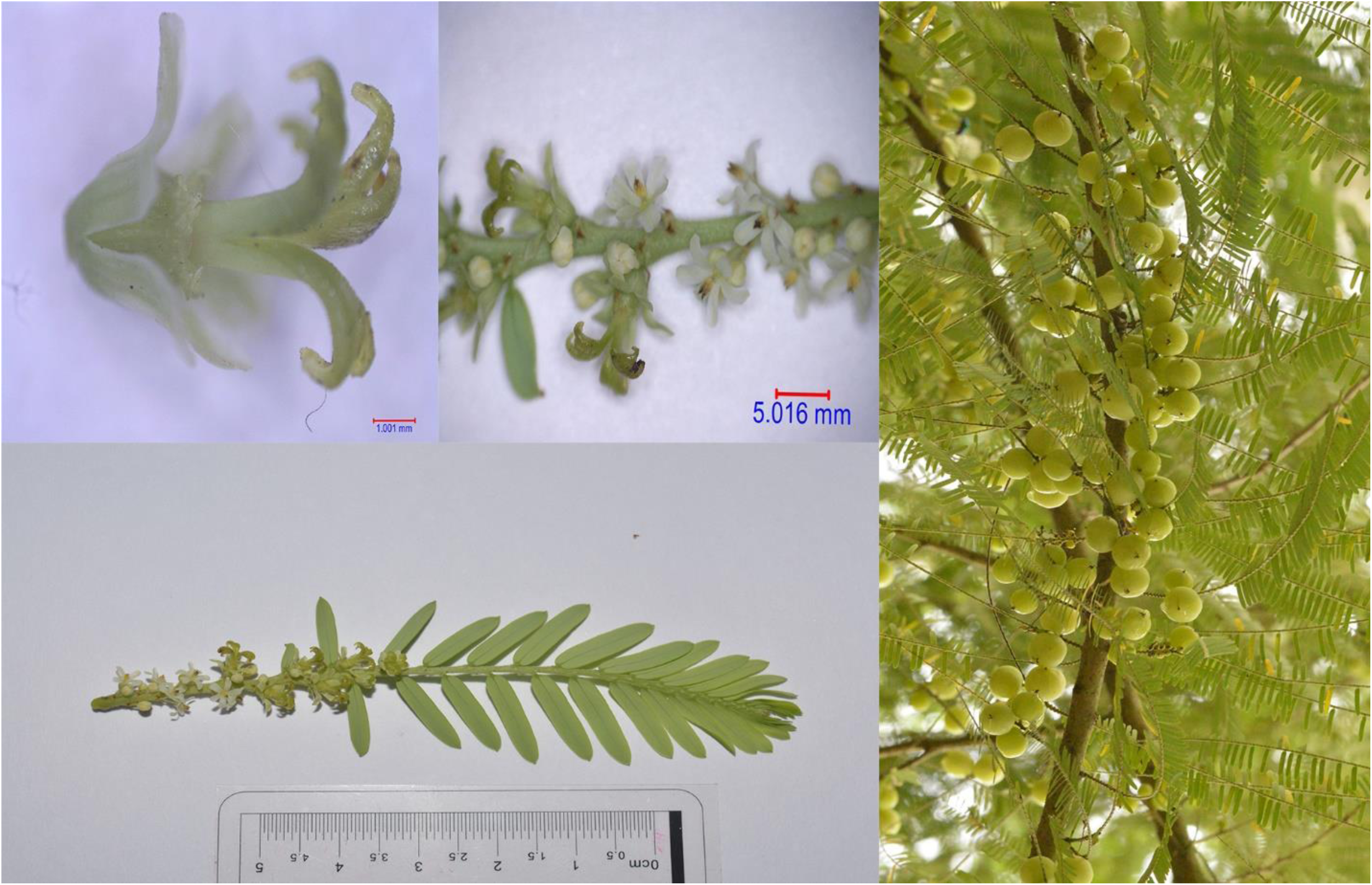
Different parts of *Phyllanthus emblica* L.

### *Phyllanthus niruri* L.: In: Sp. Pl.: 981 (1753)

#### Synonyms

*Urinaria erecta* Medik.: In: Malvenfam.: 80 (1787), *Nymphanthus niruri* (L.) Lour.: In: Fl. Cochinch.: 545 (1790), *Phyllanthus carolinianus* Blanco: In: Fl. Filip.: 691 (1837), *Niruris annua* Raf.: In: Sylva Tellur.: 91 (1838), *Niruris indica* Raf.: In: Sylva Tellur.: 91 (1838), *Phyllanthus moeroris* Oken: In: Allg. Naturgesch. 3(3): 1601 (1841), *Phyllanthus filiformis* Pav. ex Baill.: In: Recueil Observ. Bot. 1: 29 (1860), *Phyllanthus ellipticus* Buckley, nom. illeg.: In: Proc. Acad. Nat. Sci. Philadelphia 1862: 7 (1863), *Phyllanthus lathyroides* var. *commutatus* Müll.Arg.: In: Linnaea 32: 41(1863), *Phyllanthus niruri* var. *genuinus* Müll.Arg., nom. inval.: In: Prodr. 15(2): 406 (1866), *Diasperus niruri* (L.) Kuntze: In: Revis. Gen. Pl. 2: 600 (1891), *Phyllanthus lathyroides* f. decoratus Standl. & Steyerm.: In: Fieldiana, Bot. 24(6): 152 (1949), *Phyllanthus erectus* (Medik.) M.R.Almeida: In: Fl. Maharashtra 4B: 343 (2003).

A Monoecious erect annual herb, up to 70 cm high; stems often branching phyllanthoid with age, branches angular, 2-5 cm long (can reach up to 8cm), usually with 13-26 leaves. Leaves alternate, stipulate, lanceolate, scarious, acute, 1.0-1.2mm long; petiolate, petioles very short 0.3-1mm long; elliptic-obolong to elliptic-oblanceolate, 5-12 mm long and 2-5mm wide, obtuse or rounded at the apex and tapering to the base, entire leaf margin, membranous, lateral nerves 4-7 pairs, indistinct, dark green above, paler and greyish beneath. Flower bisexual, light yellow, very numerous, axillary, the male 1-3, female solitary. Male flowers are staminate, stamen 3, filamentous, filaments united into a short column; pedicellate, pedicels are 0.9-1 mm long and less than 0.5 mm width; sepal 6, 0.6-0.7 mm long, obovate, rounded, midrib yellowish green; disc gland 6, lobulated, verruculose; anther horizontal. Female are pedicellate, pedicle 1.4-1.9 mm long; sepal 6, unequal, oblong to obolanceolate, rounded, white, 1.0-1.5 mm long and 0.4-0.5 mm width, lobulate, 6 lobed; nectary disk thin and flat, deeply lobed into 6-10, some crenate and broad, some are bifid, some others linear and entire; ovary subglobose, 0.8-1 mm diameter, smooth, styles minute and free, adpressed or ascending, lobes recurved. Fruit a capsule, trilobite-subglobose, approximately 1.5-2.5 mm in diameter, the surface smooth, olivaceous or stramineous. Seeds angled ca. 1 mm long and 0.6 mm width, longitudinally 7-8 ribs on back, ochreous-fulvous, dark yellowish brown. Flowering and fruiting: August-October (Fig. 6).

**Fig. 6:**
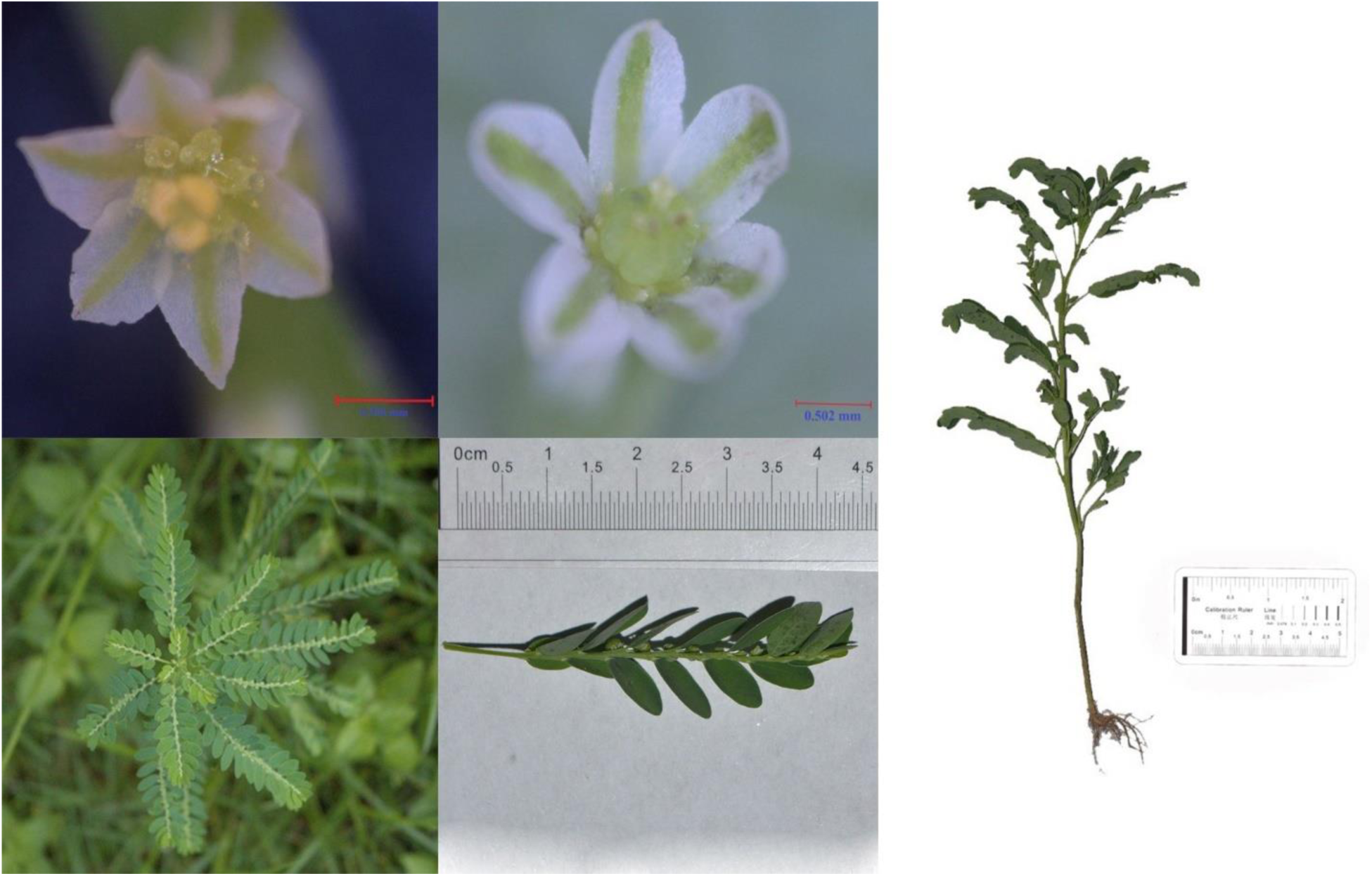
Different parts of *Phyllanthus niruri* L.

### *Phyllanthus reticulatus* Poir.: In: Encycl. 5: 298 (1804)

#### Synonyms

*Anisonema reticulatum* (Poir.) A.Juss.: In: Euphorb. Gen.: 4 (1824), *Cicca reticulata* (Poir.) Kurz: In: Forest Fl. Burma 2: 354 (1877), *Diasperus reticulatus* (Poir.) Kuntze: In: Revis. Gen. Pl. 2: 600 (1891), *Kirganelia reticulata* (Poir.) Baill.: In: Étude Euphorb.: 613 (1858), *Anisonema dubium* Blume: In: Bijdr. Fl. Ned. Ind.: 589 (1826), *Anisonema intermedium* Decne.: In: Nouv. Ann. Mus. Hist. Nat. 4: 482 (1831), *Anisonema jamaicense* (Griseb.) Griseb.: In: Fl. Brit. W. I.: 716 (1864), *Phyllanthus alaternoides* Rchb. ex Baill.: In: Recueil Observ. Bot. 1: 83 (1860), *Phyllanthus jamaicensis* Griseb.: In: Fl. Brit. W. I.: 34 (1859), *Phyllanthus sinensis* Müll.Arg.: In: Linnaea 32: 12 (1863), *Phyllanthus sinensis* var. *dalbergioides* Müll.Arg., nom. inval.: In: Linnaea 32: 12 (1863), *Phyllanthus takaoensis* Hayata: In: Icon. Pl. Formosan. 9: 94 (1920).

A Monoecious erect perineal shrub, up to 6 m high; bark peeling or flaking, grayish-brown; leaves and pedicels puberulous or plabrous, yellowish green; usually with 11-28 leaves. Leaves alternate, stipulate, stipule subulate-lanceolate, 0.8-2mm long, acuminate, truncate at base; petiolate, petioles very short 1.5-3mm long, chartaceous; leaf blade varies, mostly elliptic to ovate; 15-26 mm long and 5-15mm wide, obtuse to rounded at the apex and cuneate at the base, entire leaf margin, membranous, lateral nerves 5-9 pairs, indistinct, tertiary vines reticulate; dark green above, paler and greyish beneath. Flowers an axillary fascicle, rarely a cyme, with 2-10 male and 1 or 2 female flowers. Male flowers are staminate, stamen 5, outer 2 free, inner 3 short and united at base, longer, filamentous stout; pedicellate, pedicels are 6-10 mm long; sepal 5-6, in 2 series, ovate or obovate, unequal, entire, 0.6-1.5 mm long; disc gland 5, scale like, diameter ca. 0.5mm; anther longitidunally dehiscent, less than ca. 0.5 mm in diameter. Female are pedicellate, pedicle 3-8mm long, slender; sepal 5-6, unequal, in 2 or 3 series, oblong-elliptic or suborbicular, 1-1.5 mm long and 0.7-1.2 mm width; disk glands 5-6, free, obolong or obovate, flattened; ovary 4-12 celled, smooth, ca. 1mm; styles free, bifid at apex. Fruit a berry, globose to oblate, c 4-6 mm wide, greenish at initial stage, black and dark purplish at maturity, 4-12-celled, 8-16-seeded. Seeds trigonous, 1.6-2 mm, brownish yellow. Flowering and fruiting: March-November (Fig. 7).

**Fig. 7:**
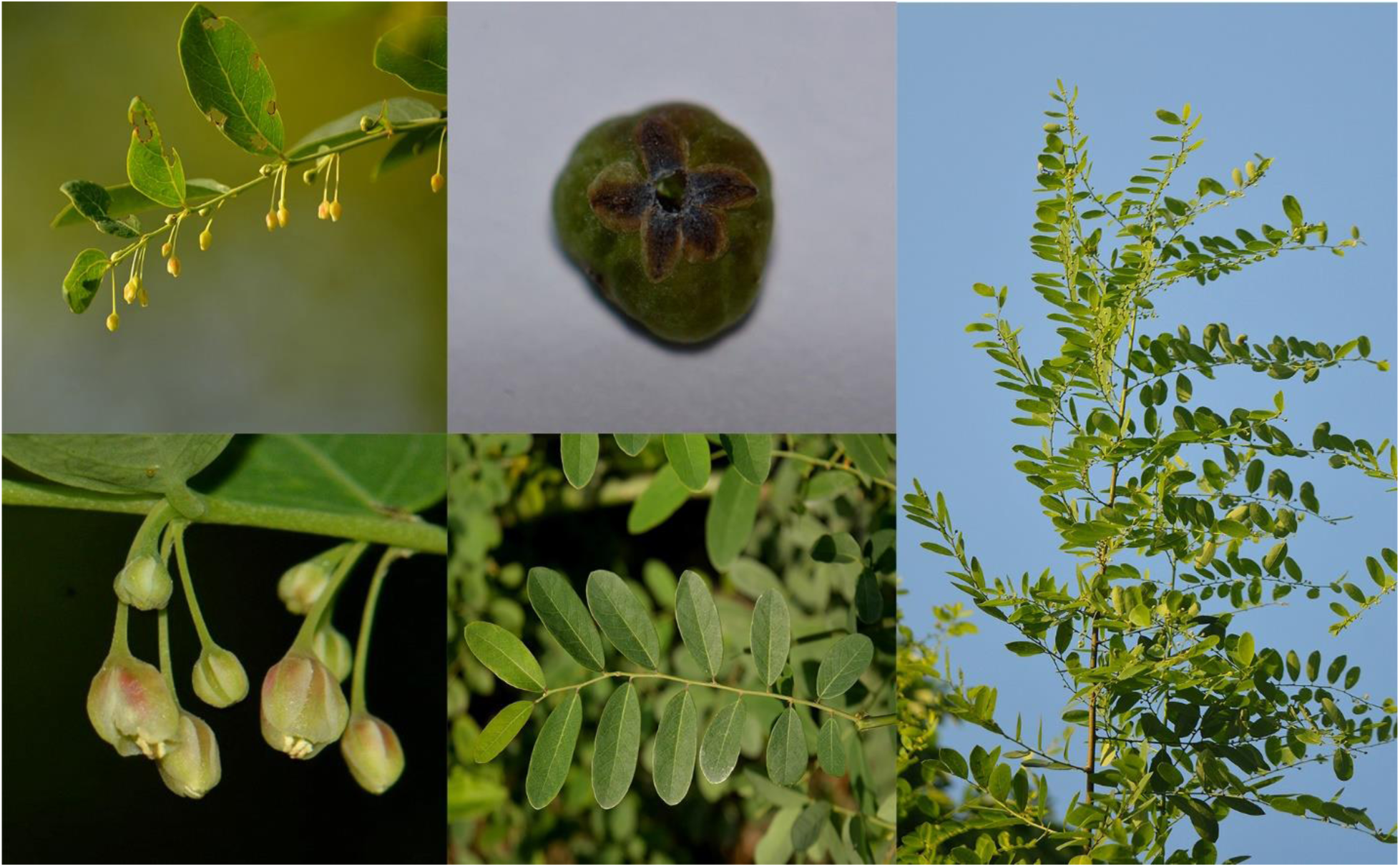
Different parts of Phyllanthus reticulatus Poir.

### *Phyllanthus urinaria* L.: In: Sp. Pl.: 982 (1753)

#### Synonyms

*Diasperus urinaria* (L.) Kuntze: In: Revis. Gen. Pl. 2: 601 (1851), *Phyllanthus alatus* Blume: In: Bijdr. Fl. Ned. Ind.: 594 (1826), *Phyllanthus cantoniensis* Hornem.: In: Enum. Pl. Hort. Hafn.: 29 (1827), *Phyllanthus croizatii* Steyerm.: In: Fieldiana, Bot. 28: 317 (1952), *Phyllanthus echinatus* Buch.-Ham. ex Benth., nom. nud.: In: Numer. List: n.° 7893B (1847), *Phyllanthus lauterbachianus* Pax: In: Repert. Spec. Nov. Regni Veg. 8: 325 (1910), *Phyllanthus leprocarpus* Wight: In: Icon. Pl. Ind. Orient. 5: t. 1895 (1852), *Phyllanthus mauritianus* Henry H.Johnst.: In: Trans. & Proc. Bot. Soc. Edinburgh 20: 329 (1895), *Phyllanthus muricatus* Benth., nom. nud.: In: Numer. List: n.° 7898D (1847), *Phyllanthus nozeranii* Rossignol & Haicour: In: Amer. J. Bot. 74: 1858 (1988), *Phyllanthus rubens* Bojer ex Baker: In: Fl. Mauritius: 309 (1877), *Phyllanthus urinaria* var. *laevis* Haines: In: Bot. Bihar Orissa 2: 125 (1921), *Phyllanthus urinaria* var. *oblongifolius* Müll.Arg.: In: Linnaea 32: 19 (1863), *Phyllanthus verrucosus* Elmer, nom. illeg.: In: Leafl. Philipp. Bot. 7: 2649 (1915).

A Monoecious erect annual herb, sometimes perineal, glabrous or puberulous, up to 1m high, tems much-branched at base, branchlets 3-12 mm long, angular, usually with 12-22 leaves. Leaves alternate, stipulate, ovate-lanceolate, 1.0-1.5mm long; petiolate, petioles very short 0.3-0.5 mm long; leaf blade linear or oblong to oblong-obovate, 5-20 mm long and 2-7 mm wide, acute apex, mucronulate, conspicuously ariculate or obtuse base; lateral nerves 4-6 pairs, conspicuous, glaucous beneath; dark greenish brown above, paler and greyish beneath. Flower bisexual, light yellow to reddish yellow, axillary, glomerules on deciduous branch. Male flowers are pedicellate, pedicels are 0.5 mm long, ariculate above the middle; staminate, stamen 3, filamentous, filaments united into a slender column; sepal 6, 0.3-0.6 mm long, elliptic to obolong-obovate, obtuse at apex; disc gland 6, green, papillose, alternate with sepal; anther erect, longitudinally dehiscent, sessile but not fused together. Female are solitary, pedicellate, pedicle 0.2-0.5 mm long; sepal 6, subequal, oblong-lanceolate, obtuse or subacute, subglabrous, yellowish with a reddish-olive midrib, 0.7-1.1 mm long and 0.2-0.5 mm width; nectary disk flat, entire; styles 3, minute and free, adpressed to the top of the ovary; ovary bifid, lobes recurved. Fruit a capsule, globose, 2-2.5 mm in diameter, with reddish blotches, scurfy-tuberculate. Seed 3-sided, 1-1.2mm long and 0.9-1 mm wide, light grayish brown, with 12-15 sharp transverse ridges on back and sides, often with 1-3 deep circular pits on side. Flowering and fruiting: April-October (Fig. 8).

**Fig. 8:**
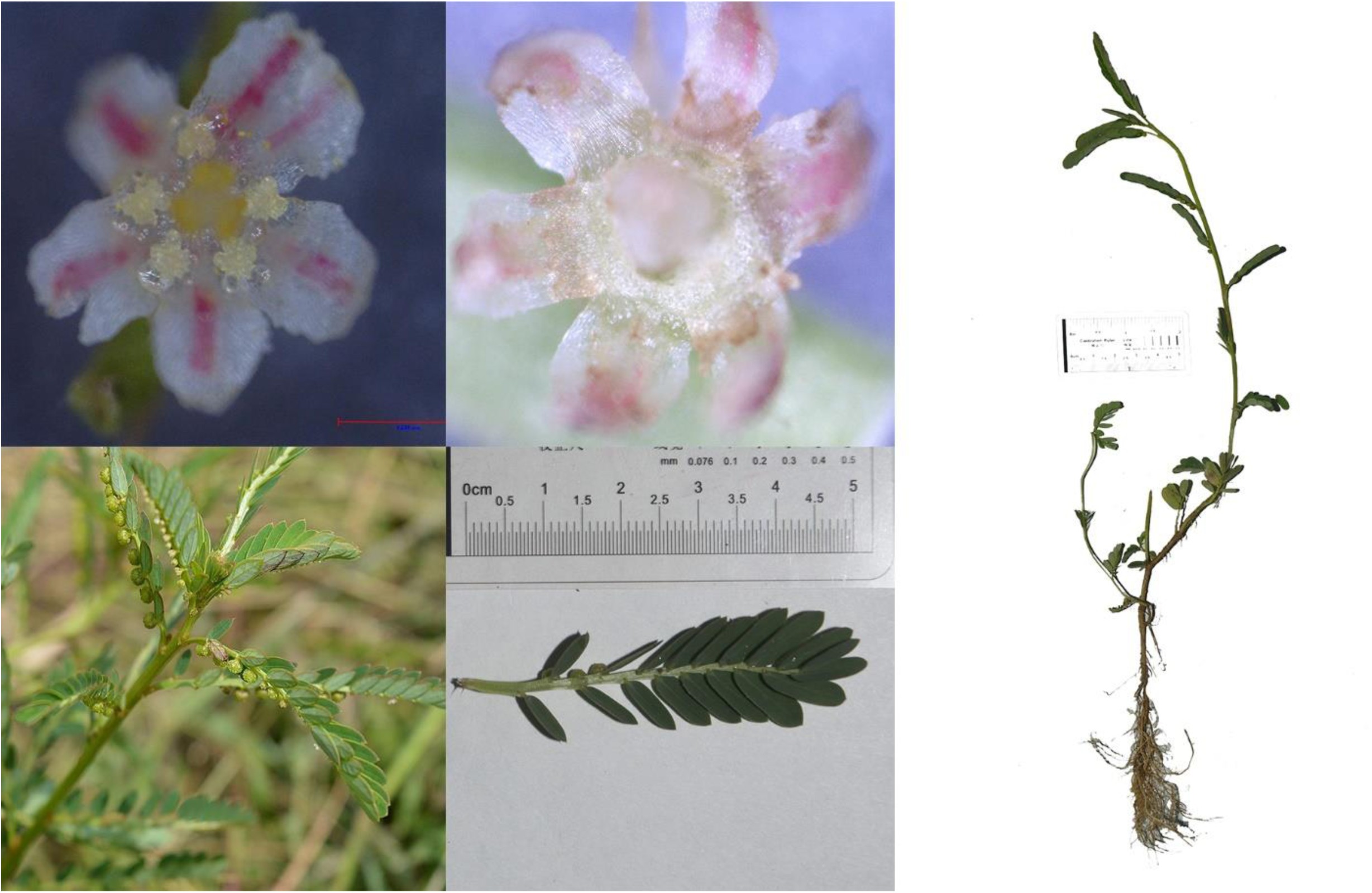
Different parts of *Phyllanthus urinaria* L.

### *Phyllanthus virgatus* G.Forst.: In: Fl. Ins. Austr.: 65 (1786)

#### Synonyms

*Phyllanthus simplex* var. *virgatus* (G.Forst.) Müll.Arg., nom. illeg.: In: Linnaea 32: 32 (1863), *Diasperus virgatus* (G.Forst.) Kuntze: In: Revis. Gen. Pl. 2: 597 (1891), *Diasperus beckleri* (Müll.Arg.) Kuntze: In: Revis. Gen. Pl. 2: 598 (1891), *Diasperus conterminus* (Müll.Arg.) Kuntze: In: Revis. Gen. Pl. 2: 599 (1891), *Diasperus depressus* Kuntze: In: Revis. Gen. Pl. 2: 599 (1891), *Diasperus minutiflorus* (F.Muell. ex Müll.Arg.) Kuntze: In: Revis. Gen. Pl. 2: 600 (1891), *Diasperus miquelianus* (Müll.Arg.) Kuntze: In: Revis. Gen. Pl. 2: 600 (1891), *Diasperus pedunculatus* (Kostel.) Kuntze: In: Revis. Gen. Pl. 2: 597 (1891), *Macraea oblongifolia* Wight: In: Icon. Pl. Ind. Orient. 5(2): 27 (1852), *Melanthesa anceps* (Vahl) Miq.: In: Fl. Ned. Ind. 1(2): 371 (1859), *Phyllanthus anceps* Vahl: In: Symb. Bot. 2: 95 (1791), *Phyllanthus beckleri* Müll.Arg.: In: Linnaea 34: 74 (1865), *Phyllanthus eboracensis* S.Moore: In: J. Linn. Soc., Bot. 45: 216 (1920), *Phyllanthus patens* Miq. ex Müll.Arg.: In: Linnaea 32: 34 (1863), *Phyllanthus simplex* Retz.: In: Observ. Bot. 5: 29 (1789), *Phyllanthus simplex* var. *myrtifolius* Domin: In: Biblioth. Bot. 22: 876 (1927), *Phyllanthus weinlandii* K.Schum.: In: Fl. Schutzgeb. Südsee, Nachtr.: 287 (1905).

A monoecious erect annual herb, sometimes perineal, up to 70 cm high, glabrous throughout, branchlets angled, more or less compressed, smooth; stem usually slightly woody at base. Leaves stipulate, stipule 0.8-1.0mm long, membranous, ovate, acuminate at apex, prominently and asymmetrically auriculate at base; petiolate, petioles very short 0.6-0.9 mm long; leaf blade linear-lanceolate, oblong or narrowly elliptic, 5-18 mm long and 2-8mm wide, obtuse or acute at the apex, rounded and slightly oblique at the base, entire leaf margin, membranous, lateral nerves obscure at both surface. Flower bisexual, axillary fascicled, usually 2-4 male and one female per fascicle.Male flowers are pedicellate, pedicels are 0.9-1 mm long; sepal 6, 0.4-0.6 mm long, broadly elliptic, ovate or orbicular, obolong; staminate, stamen 3, filamentous, filaments are 0.3-0.5 mm long; disc gland 6, lobulated; anther subglobose, 0.2-0.3 mm in diameter, horizontally dehiscent. Female are pedicellate, pedicle 3-5 mm long; sepal 6, subequal, ovate-oblong, acute to obtuse, margin slightly white-membranous, reflected in fruits, purple, 0.7-1.0 mm long, lobulate, 6 lobed; nectary disk thin and flat, orbicular, entire or subentire, ca. 0.6 mm diameter; ovary globose, 0.2-3 mm diameter,, bifid; smooth, styles minute and free, 3-celled, ca. 0.3 mm long, revolute. Fruiting occurred at pedicels, fruiting pedicels are 5-12 mm long; fruit is a capsule, capsules oblate, 2-3 mm in diameter, purple, with raised scales or smooth. Seeds trigonous, 1.2-1.5 mm, finely warty. Flowering and fruiting: May-October (Fig. 9).

**Fig 9:**
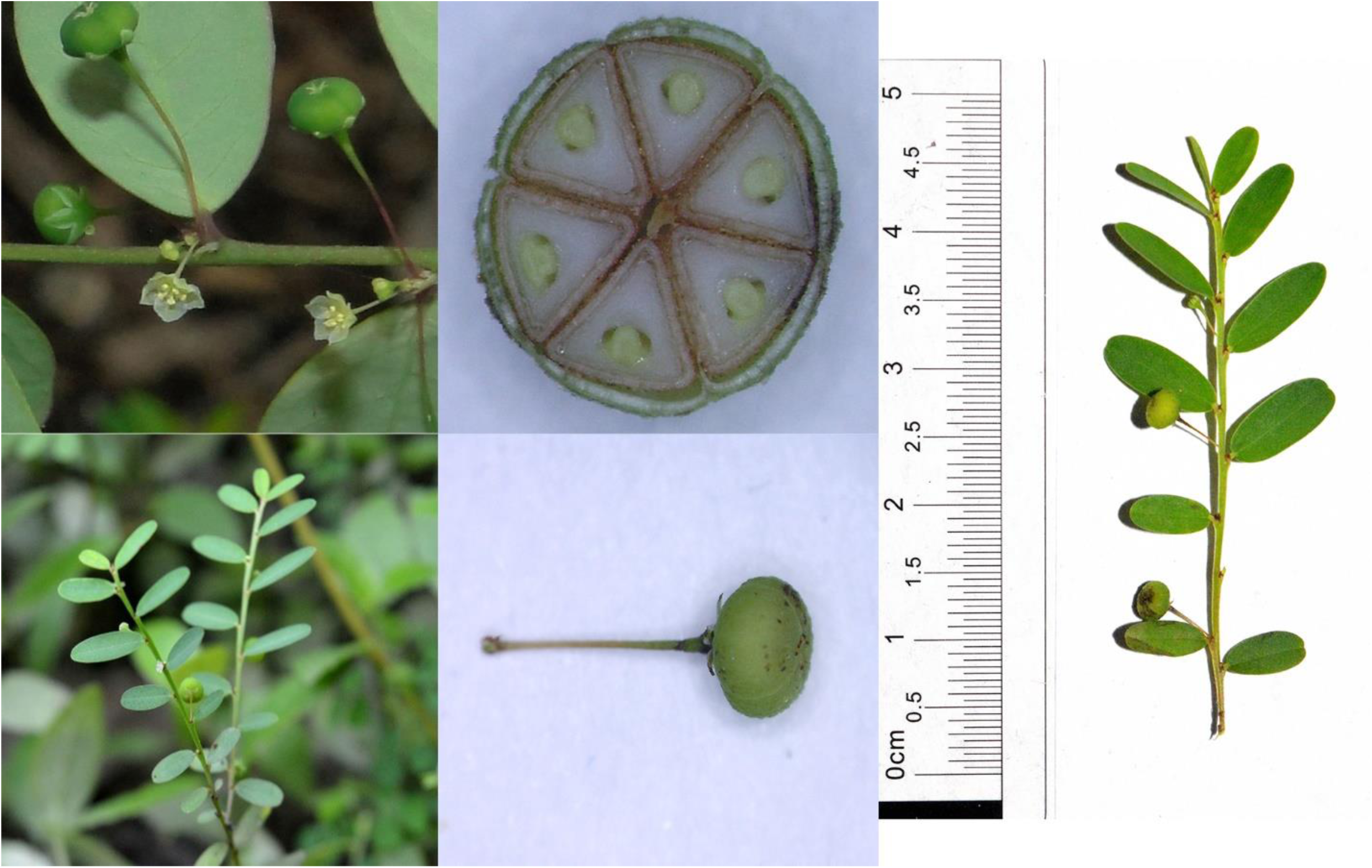
Different parts of *Phyllanthus virgatus* G.Forst.

### Key to identify the Phyllanthus of South-western Bangladesh

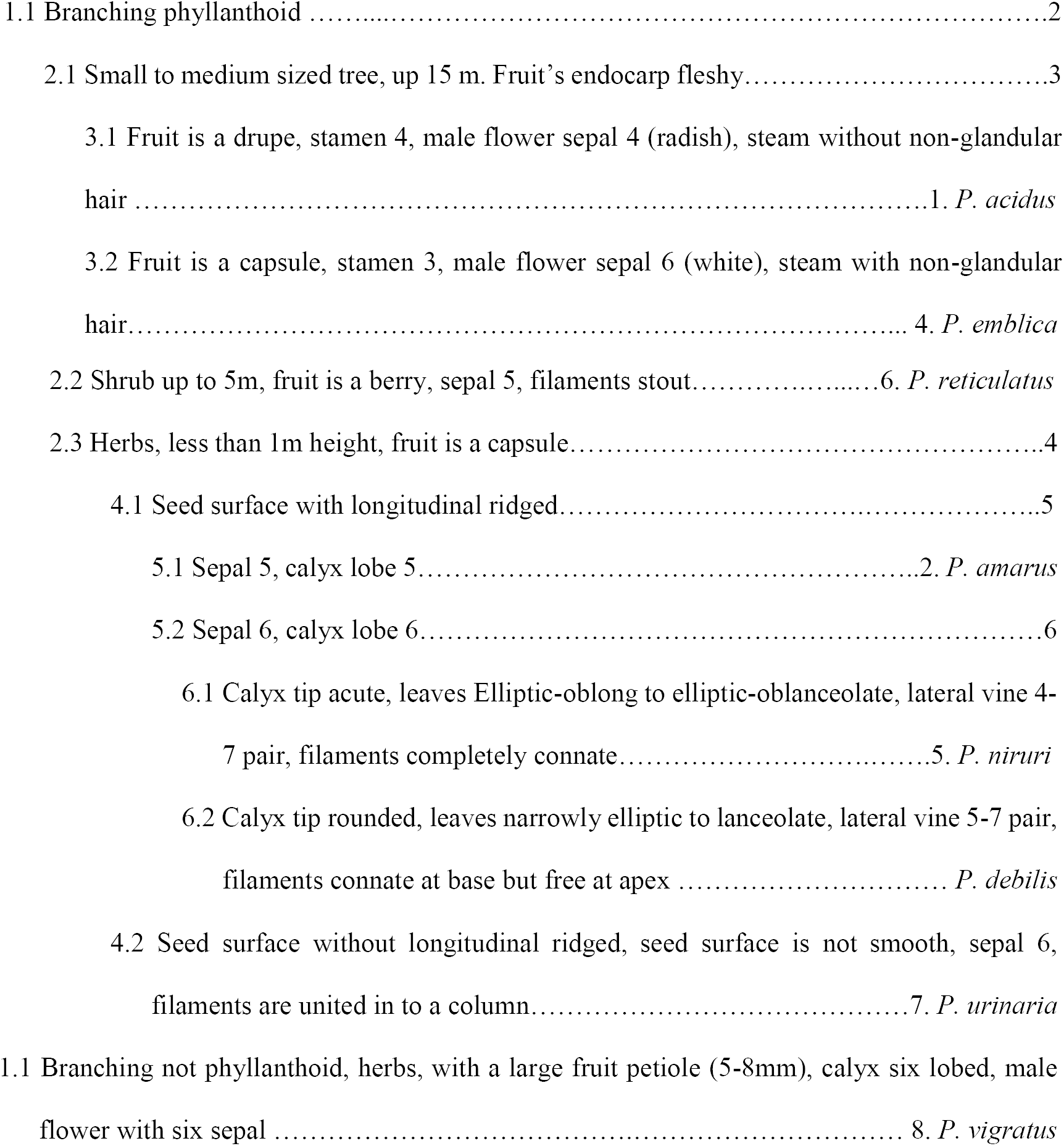

### Anatomical observation

The epidermis of the studied eight species was one to three layer. The mesophylls are differentiated into palisade and spongy parenchyma layer but those varied from species to species. Among the studied species, epidermal circumference found rounded for *P. amarus, P. reticulatus, P. acidus*, and angular for *P. emblica, P. debilis, P. niruri*, and *P. urinaria*. *P. vigratus* was the only species which had the elliptical circumference only. Besides, *P. urinaria* had a special anatomic feature called rings and farrows were observed in cortex. In consideration of hypodermis, most of the species found rectangular except *P. vigratus* had a thick oval shape hypodermis composed of 3-6 layers. Structure of the studied species cortex had a wide range of variations. *P. amarus, P. niruri*, P. urinaria, *P. reticulatus* and *P. acidus* had oval shaped cortex. On the other hand, *P. emblica* had a sub-oval shaped cortex, elliptical for *P. vigratus* and rectangular for *P. debilis*. All the species found a moderately thick layered parenchyma composed of 1-15 layer parenchyma or spongy parenchyma cells. *P. acidus* had the thickest layer of parenchyma cells which was 8-15 layered thick (9 A). For all of the studied species, phloem section was thicker than xylem. *P. acidus* had the thickest layer of phloem cells which was composed of 5-15 layers (table 4, Fig 10).

**Table 3:** Morphological comparisons among studied *Phyllanthus*.

|  | *P. amarus* | *P. debilis* | *P. niruri* | *P. urinaria* | *P. virgatus* | *P. reticulatus* | *P. acidus* | *P. emblica* |
| --- | --- | --- | --- | --- | --- | --- | --- | --- |
| **Form** | Herb | Herb | Herb | Herb | Herb | Shrub | Tree | Tree |
| **Branchlets** | Present | Present | Present | Present | Absent | Present | Present | Present |
| **Leaf shape** | Oblong or elliptic-oblong | Narrowly elliptic to lanceolate | Elliptic-oblong to elliptic-ob lanceolate | Linear-oblong to oblong-obovate | Obolong-elliptic | Varying but mostly obtuse to rounded | Ovate to ovate-lanceolate | Oblong or linear-oblong |
| **Leaf apex** | Cbtuse or rounded | Acute to acuminate | Obtuse or rounded | Acute | Obtuse or acuminate | Obtuse to rounded | Acute to acuminate | Obtuse |
| **Leaf base** | Rounded | Cuneate to acute | Tapering | Mucronulate, conspicuously ariculate or obtuse | prominently and asymmetrically auriculate | Cuneate | Rounded or broadly cuneate. | Shallowly cordate and slightly oblique |
| **Lateral vine** | 4-7 pair | 5-7 pair | 4-7 pair | 4-6 pair | 6-9 pair | 5-9 pair | 5-7 pair | 4-9 pair |
| **Sepals** | Five | Six | Six | Six | Six | Five | Four | Six |
| **Disk segments** | Five | Six | Six | Six | Six | Five | Four | Six |
| **Filaments** | Completely connate | Connate at base and free at apex | Completely connate | Connate, united into a slender column | Connate, Free | Stout | Kidney shaped. | Stout |
| **Style** | Three, Minutely bifid at tip. | Three, Minutely bifid at middle. | Three, triangular and bifid. | Three, free and adpressed to the top of ovary. | Three, boiled at apex | Five, two free and three united. | Four, free and two shorter. | Two parted |
| **Fruit** | Capsule | Capsule | Capsule | Capsule | Capsule | Barry | Drupe | Capsule |
| **Calyx** | Five lobed | Six lobed (apex rounded) | Six lobed (apex acute) | Six lobed | Six lobed | Five lobed | Six lobed | Six lobed |

**Table 4:** Anatomical comparison of stem cross section among studied *Phyllanthus*.

|  | *P. amarus* | *P. debilis* | *P. niruri* | *P. urinaria* | *P. virgatus* | *P. reticulatus* | *P. acidus* | *P. emblica* |
| --- | --- | --- | --- | --- | --- | --- | --- | --- |
| **Epidermal circumference** | Rounded | Angular | Angular | Angular | Elliptical | Rounded | Rounded | Rounded |
| **Ring and Farrows** | Absent | Absent | Absent | Present | Absent | Absent | Absent | Absent |
| **Hypodermis** | Rectangular<br>2-3 layer<br>thick | Rectangular<br>4-5 layer<br>thick | Rectangular<br>2-3 layer<br>thick | Rectangular<br>2-5 layer<br>thick | Rectangular<br>3-4 layer<br>thick | Rectangular<br>3-5 layer<br>thick | Rectangu<br>lar<br>2-5 layer<br>thick | Oval<br>3-6 layer thick |
| **Cortex** | Oval | Rectangular | Oval | Oval | Elliptical | Oval | Oval | Sub-oval |
| **Parenchyma** | 4-6 layer<br>thick | 3-5 layer<br>thick | 3-6 layer<br>thick | 2-5 layer<br>thick | 1-3 layer<br>thick | 3-5 layer<br>thick | 8-15<br>layer<br>thick | 6-8 layer thick |
| **Pericycle** | 2-3 layer<br>thick | 3-5 layer<br>thick | 3-5 layer<br>thick | 2-4 layer<br>thick | 2-3 layer<br>thick | 2-7 layer<br>thick | 2-8 layer<br>thick | 2-5 layer thick |
| **Xylem** | 3-6 layer<br>thick | 1-3 layer<br>thick | 2-4 layer<br>thick | 2-5 layer<br>thick | 1-3 layer<br>thick | 2-4 layer<br>thick | 1-3 layer<br>thick | 1-3 layer thick |
| **Phloem** | 4-9 layer<br>thick | 5-8 layer<br>thick | 4-7 layer<br>thick | 3-5 layer<br>thick | 2-3 layer<br>thick | 4-7 layer<br>thick | 5-15<br>layer<br>thick | 5-8 layer thick |

**Fig 10:**
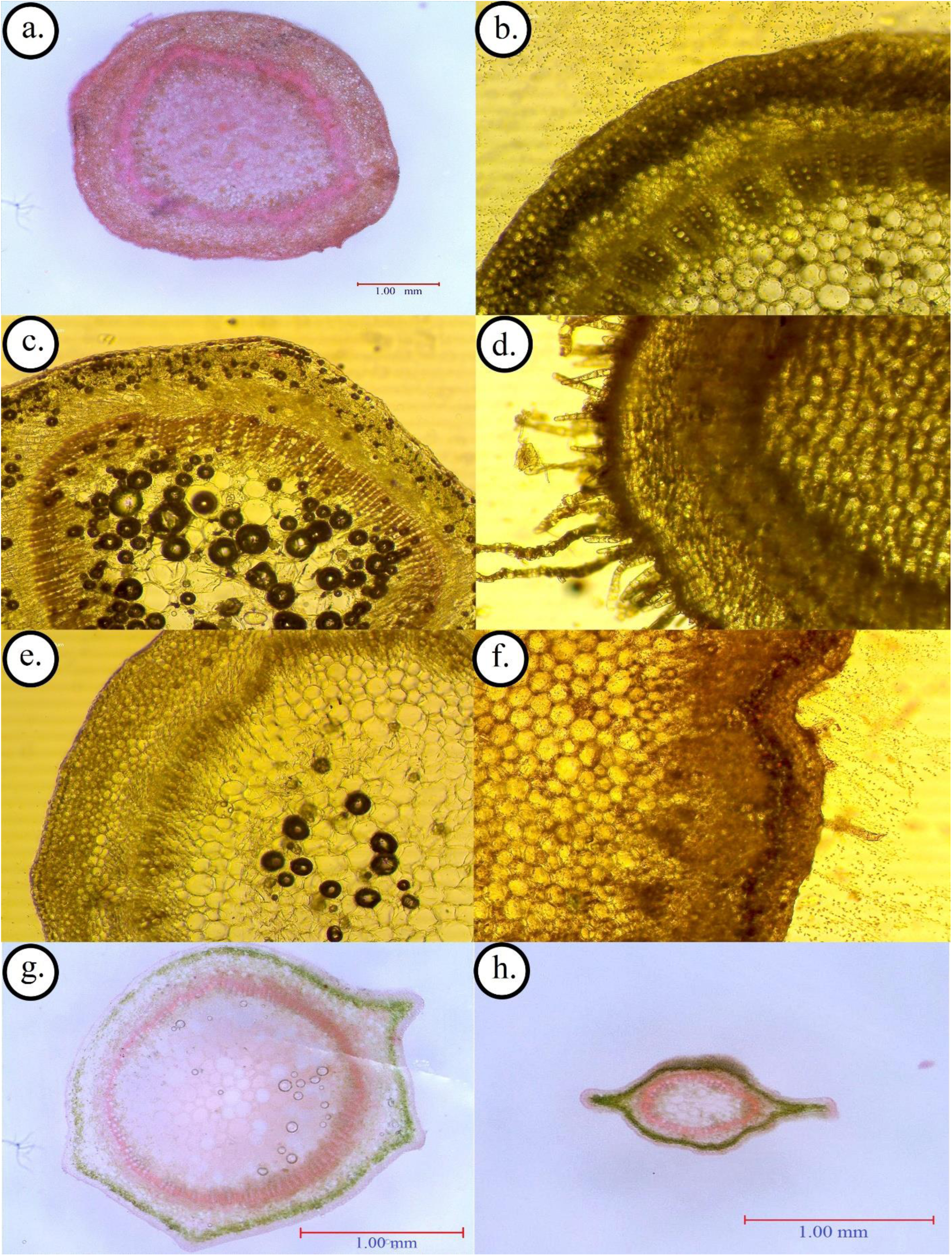
Cross section of stem of phyllanthus of southwestern Bangladesh. (a: *P. acidus*, b: *P. amarus*, c: *P. debilis* d: *P. emblica*, e: *P. niruri*, f: *P. reticulatus*, g: *P. urinaria*, h: *P. vigratus*)

## Discussion

In Bangladesh, very limited works were found that addressing to identify Phyllanthus sp. Ahamed et al. (2008) reported 11 species representing Phyllanthus genus in Bangladesh including 5 herbs, 4 shrubs and 2 trees. This study found 5 herbs, 1 shrubs and 2 trees in the study area including two new species (*Phyllanthus debilis* and *Phyllanthus ama*rus) for Bangladeshi flora. Besides application of anatomical features along with taxonomic notes improve the identification of species (Kandavel et al. 2011).

Among listed five Phyllanthus herbs, *P. urinaria* and *P. virgatus* were easy to differentiate than others, based on morphological and anatomical features. With rough fruit surface for P. *urinaria* and long fruit petiole for *P. virgatus*, they were conveniently identifiable in field surveys. *P. amarus, P. debilis* and *P. niruri* were morphologically very close and difficult to differentiate. With five calyx segments *P. amarus* was different than *P. niruri* and *P. debilis* with six calyx segments. In the current study, filament stricture found most convenient option to differentiate among them *P. niruri* observed completely connate and joined filaments but for *P. debilis* they were connate at base but free at apex (table 3). In terms of anatomy, *P. amarus* have rounded epidermal circumference but *P. niruri* and *P. debilis* had angular epidermal circumference. Hypodermis layer found key features for differentiate among P. niruri and *P. debilis*. More than 3 layered rectangular cell *P. debilis* Hypodermis was different than *P. niruri*, which has less than or exact 3 layered rectangular hypodermis (table 4; Fig 10).

Both *Phyllanthus debilis* and *Phyllanthus ama*rus were first recorded from India by J. D. Hooker (1890) from North-west India, Sikkim, Bhutan, Hazaribag, Behar and Assam (Hooker, 1890). Prain (1904) also recorded them from Orissa, Chota Nagpur, Behar, Trihut and North Bengal (Prain 1904). In recent days these species were listed in Chennai, Tamil Nadu; Great Nicobar (Srirama et al. 2012). However, they were absent in all regional floristic works of Bangladesh (Rahman & Jamila 2016; Kona & Rahman 2015; Rahman et al. 2015; Rahman et al. 2014; Rahman 2013; Rahman & Akter 2013; Rahman et al. 2013; Uddin & Hassan 2012; Uddin et al. 2013; Arefin et al. 2011; Uddin & Hassan 2010; Tutul et al. 2010; Islam et al. 2009; Ahamed et al. 2008; Hossain et al. 2005; Khan & Huq 2001; Mia & Khan 1995; Datta & Mitra 1953). These facts, ensure that this two species were new to the floral database of Bangladesh. This finding will help the path of ecological study of this species for their conservation in Bangladesh.

A specimen *Phyllanthus debilis* J. G. Klein ex Willd.a was collected from Khulna University Campus ((22°48′01.89″N; 89°32′04.60″E) in late October 2015. This species was found in a built-up habitat of Khulna University Nursery and was associated with *Phyllanthus niruri* L., *Hedyotis corymbosa* (L.) Lam., *Cynodon dactylon* (L.) Pers., *Cyperus rotundus* L., *Synedrella nodiflora* (L.) Gaertn., *Scoparia dulcis* L., *Lindernia crustacea* (L.) F. Muell., *Imperata cylindrical* (L.) Raeusch, *Euphorbia hirta* L. and others. Specimen of *Phyllanthus amarus* Schumach. & Thonn. was collected from Jessore city area (23° 09′55.97″N; 89°12′15.27″E) in late April of 2017. It was in association with *Synedrella nodiflora* (L.) Gaertn., *Scoparia dulcis* L., *Phyllanthus niruri* L., *Hedyotis corymbosa* (L.) Lam., *Cyperus rotundus* L., *Lindernia crustacea* (L.) F. Muell., *Euphorbia thymifolia* L., *Euphorbia microphylla* Lam. Both specimens were identified with key provided by Hooker (1890), Prain (1904) and Wurdack et al. (2004).

## Acknowledgements

Authors are thankful to Prof. A.K. Fazlul Hoque for his comments on the initial version of the manuscript. They also acknowledge Prof. Dr. Iftekhar Shams for lab support and encouragement for this study.

